# Fast and memory-efficient noisy read overlapping with KD-trees

**DOI:** 10.1101/166835

**Authors:** Dmitri Parkhomchuk, Andreas Bremges, Alice C. McHardy

## Abstract

**Motivation:** Third-generation sequencing technologies produce long, but noisy reads with increasing sequencing throughput and decreasing per-base costs. Detecting read-to-read overlaps in such data is the most computationally intensive step in *de novo* assembly. Recently, efficient algorithms were developed for this task; nearly all of these utilize long *k*-mers (>10 bp) to compare reads, but vary in their approaches to indexing, hashing, filtering, and dimensionality reduction.

**Results:** We describe an algorithm for efficient overlap detection that directly compares the full spectrum of short *k*-mers, namely tetramers, through geometric embedding and approximate nearest neighbor search in multidimensional KD-trees. A proof of concept implementation detected read-to-read overlaps in bacterial PacBio and ONT datasets with notably lower memory consumption than state-of-the-art approaches and allowed downstream *de novo* assembly into single contigs. We also introduce a sequence-context dependent tagging scheme that contributes to memory and computational efficiency and could be used with other aligning and overlapping algorithms.

**Availability:** A C++14 implementation is available under the open source Apache License 2.0 at: https://github.com/dzif/kd-tree-overlapper

## 1 Introduction

Genome sequencing and assembly of cultured microbial isolates turned from a challenge into a routine, primarily due to the advent of long-read sequencing (Goodwin *et al*., 2016; Wibberg *et al*., 2016). Technologies offered by Pacific Biosciences (PacBio; Menlo Park, CA, USA) and Oxford Nanopore Technologies (ONT; Oxford, UK) currently generate multi-kb sequencing reads resolving long genomic repeats, but deliver a high per-base error rate of around 16% (Laehnemann *et al*., 2016). This inherent noise complicates the identification of read-to-read overlaps, which is typically a prerequisite for *de novo* Overlap-Layout-Consensus (OLC) assembly (Berlin *et al*., 2015; Loman *et al*., 2015).

A read-to-read overlap occurs when two reads originate from overlapping genomic regions and thus share the same (local) sequence. General-purpose alignment algorithms have a quadratic dependence on the number of reads when performing an all-vs-all pairwise alignments and thus do not scale well. Moreover, the full base-to-base alignment is not needed to construct an overlap graph; it is sufficient to indicate read-to-read overlap candidates together with their relative orientation (Chu *et al*., 2017).

As a consequence, multiple programs were developed to efficiently overlap long noisy reads, such as BLASR (Chin *et al*., 2013), DALIGNER (Myers, 2014), MHAP (Berlin *et al*., 2015), GraphMap (Sović *et al*., 2016), and Minimap (Li, 2016). These all search for shared seeds between reads, but differ in the way these seeds are found and thereafter used to determine overlap candidates. A recent review highlights algorithmic features of the available software and evaluates their performance (Chu *et al*., 2017).

Here, we describe an algorithm that efficiently determines read-to-read overlap candidates by directly comparing short *k*-mer (tetramer) spectra of reads. It maintains a low memory footprint using geometric embedding (*k*mer counting) and approximate nearest neighbor (ANN) search (Muja and Lowe, 2014) in a KD-tree index (Bentley, 1975). We show that this approach is as precise as other methods and allows the subsequent assembly of MB-sized bacterial genomes into single contigs.

## 2 Methods

The algorithm consists of five steps: (1) tagging of long reads, to form a set of read subsequences of fixed length; (2) geometric embedding (*k*-mer counting) of each tag; (3) KD-tree index creation; (4) searching approximate nearest neighbors (ANNs) for each tag in this index; and (5) filtering for read overlaps. All steps were implemented in a C++14 program available at https://github.com/dzif/kd-tree-overlapper. For convenience, we also provide a statically linked binary and a Docker container adopting the bioboxes standard (Belmann *et al*., 2015).

### 2.1 Tagging of long reads

In this first step, sequence tags, i.e. subsequences of fixed-length, are placed on each read, because eventually a comparison of all tetramer counts between sequences will be utilized (which can straightforwardly be done for tags of the same length). In principle, read tagging can be done by covering long reads either regularly or randomly with the desired tags density. A simple algorithm of GC-profile peak detection is used to anchor tags, because the GC-profile is relatively robust against sequencing errors and invariant to reverse-complementation.

Per default, the GC content is calculated for a sliding window of 100 bp and tags of length 1200 bp are placed at the GC maxima, only allowing the distance between tags to be above a 400 bp. This places tags non-randomly and decreases the overall tag density while keeping all tags aligned in true overlaps, due to them sharing GC-profile peaks in the same subsequences relating to the underlying genome sequence (Fig. 1). This also prevents that the number of unique tags to grow linearly with increasing sequencing coverage and helps to keep the index size small.

**Figure 1:**
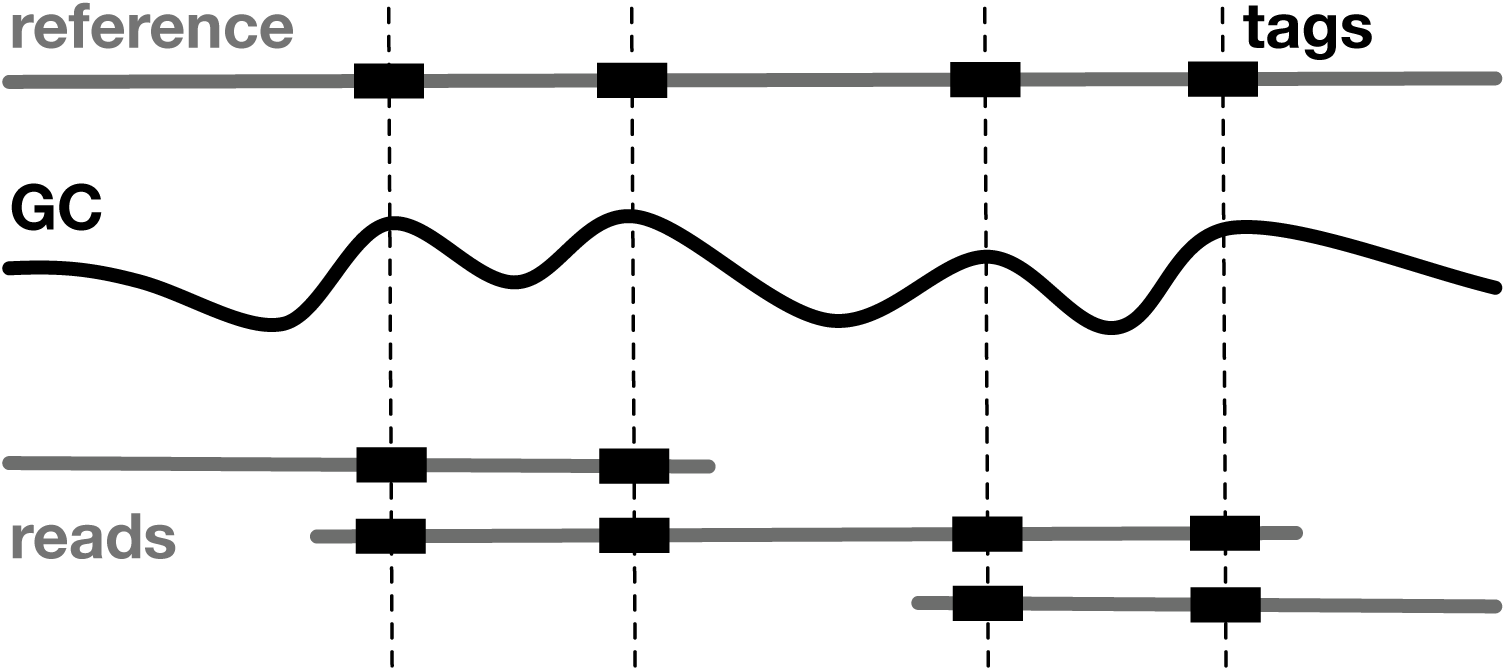
Tagging of long reads. Sequence tags, i.e. subsequences of fixed-length, are anchored at GC-profile peaks and thus aligned in true overlaps. Shared tags between different reads indicate read-to-read overlap candidates and their relative orientation can be determined if they share at least two pairs of tags.

### 2.2 Geometric embedding of each tag

Geometric embedding, or *k*-mer counting, maps a sequence (here: a tag) to a vector of counts (VC) of *k*-mers; the VC contains the ordered set of *k*-mer occurences in a sequence (including zero counts). In our implementation, *k*-mers and their reverse-complements are collapsed in the VC (Figure 2). If two tags show substantial overlap and have similar sequences the distance between their VCs will be small, corresponding to a small edit distance between the respective tags. Hence, searching for the nearest neighbors of a tag’s VC will reveal tags with high sequence similarity to the query tag.

**Figure 2:**
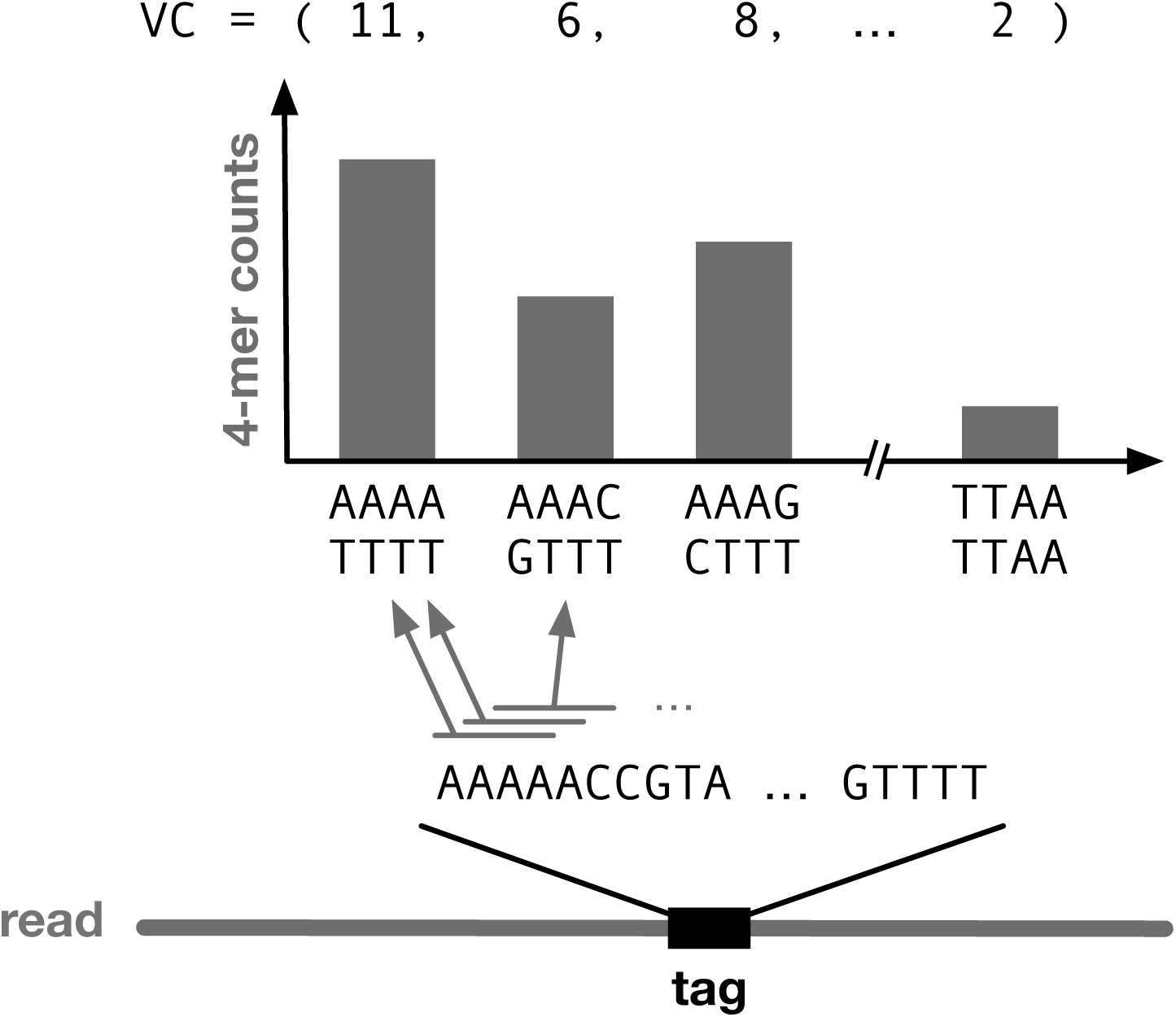
Geometric embedding of each tag. A sequence tag is mapped to a vector of counts (VC) of canonical *k*-mers, namely tetramers. A small edit distance between two tags also causes a small distance between their VCs, thus enabling an efficient detection of read-to-read overlap candidates.

### 2.3 KD-tree index creation

To identify similar VCs (and thus read-to-read overlap candidates) efficiently, we first build a KD-tree index of all VCs using the FLANN software library (Muja and Lowe, 2014). KD-trees are a generalization of binary search trees for multiple dimensions and allow an efficient exact search in low-dimensional data (Bentley, 1975). Heuristic methods for finding approximate nearest neighbors (ANNs) in high-dimensional data accelerate the search by several orders of magnitude (Muja and Lowe, 2014).

### 2.4 Approximate nearest neighbor search

An exact search in high-dimensional KD-trees is inefficient because of the dimensionality curse (Marimont and Shapiro, 1979); roughly speaking, the search space grows exponentially with the number of dimensions. However, searching for approximate nearest neighbors (ANNs) is several orders of magnitude faster than exact KD-tree query methods (Muja and Lowe, 2014). The speed-up is particularly evident for natural, high-dimensional data with high redundancy (i.e. correlation) among the input features, e.g. the number of ‘AAAA’ tetramers in a sequence is usually positively correlated with the number of ‘AAAC’ tetramers because they overlap (Figure 2). ANNs for all VCs are searched for using the KD-tree index created in the previous step.

### 2.5 Filtering of read overlaps

At this point, each VC (and thus each tag) has been assigned a list of ANNs. ANNs of tags found on different reads indicate that these originate from the same underlying sequence. Double (or more) hits, corresponding to two pairs of tags being nearest neighbors to each other and with the tags of each pair being spread across the same two reads, are extracted. Read-to-read overlap candidates are reported based on the orientation of the tag pairs relative to each other in their correct orientation (Figure 1). They are reported in the MHAP output format, a tabular format compatible with e.g. the Canu assembler (Koren *et al*., 2017).

The conceptual correspondence between the overlapping read pairs and the KD-tree index is shown in Figure 3.

**Figure 3:**
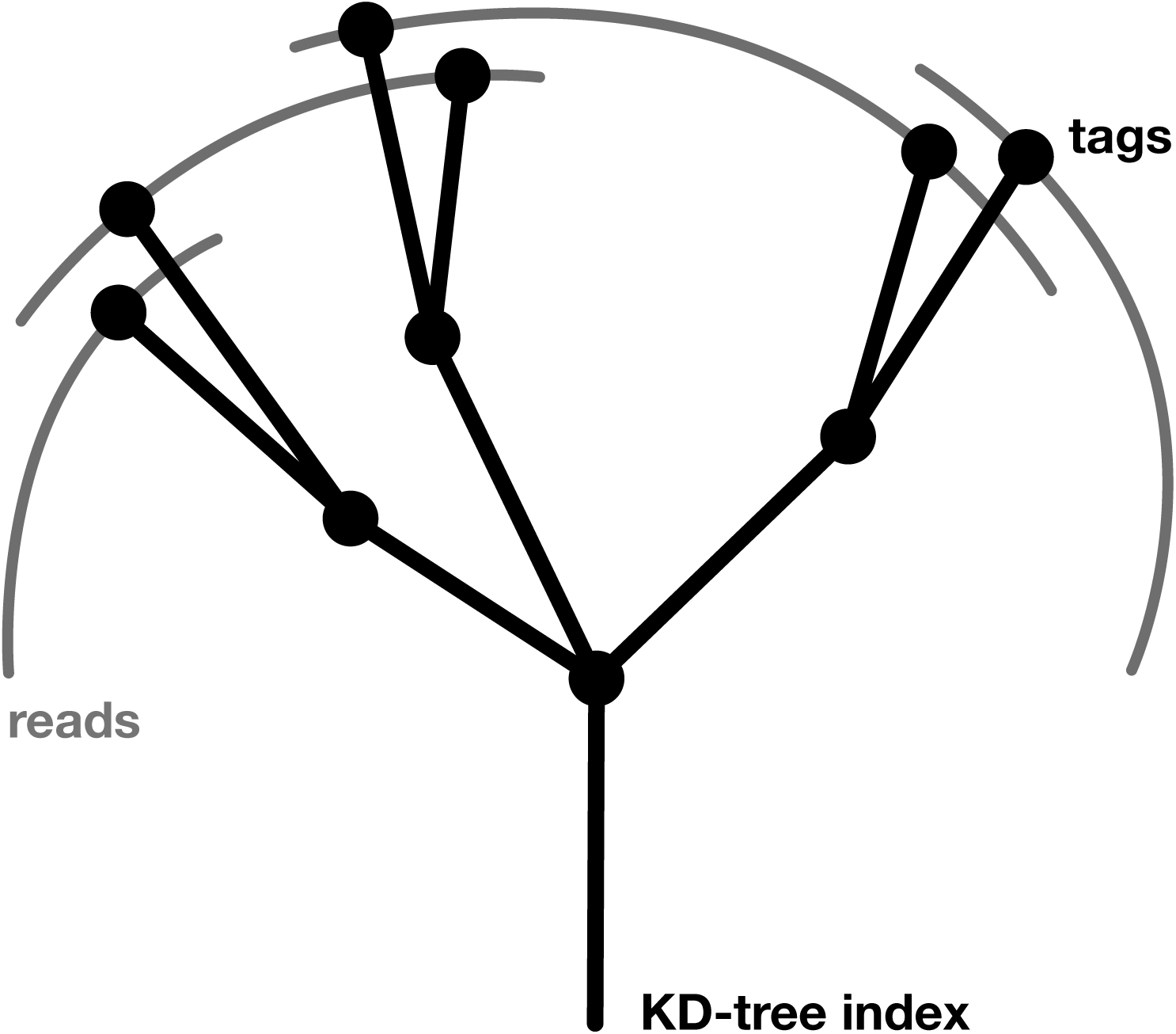
Schematic representation of overlapping reads and the KD-tree index. Shared tags between two reads indicate read-to-read overlap candidates. These tags’s VCs are located close-by in the KD-tree and are detected efficiently using approximate nearest neighbors (ANNs) search.

## 3 Results

Building upon the recent evaluation of long read overlappers, we benchmarked KD-tree using the datasets from Chu *et al*., 2017, i.e. ONT SQKMAP-006 and PacBio P6-C4 *E. coli* sequencing reads. We include their results, representing the “best” setting for each tool (those with the highest F1 score after parameter optimization), and additionally ran MHAP v2.1.1, Minimap v0.2-r124, and KD-tree (kd) v1.0 with default and recommended settings. All evaluation steps are documented in a GitHub repository (https://github.com/dzif/kd-tree-evaluation) to enable computational reproducibility.

### 3.1 Sensitivity and precision

We experimented with *k*-mers of up to *k*=7 on simulated and real long read datasets to empirically determine the *k*-mer length providing the best performance in terms of sensitivity, precision, CPU consumption and RAM requirements for KD-tree. We found that trimer spectra are not specific enough to detect true overlaps with a high precision and observed that sometimes, unrelated sequences had similar VCs by chance (data not shown). However, in tests with simulated reads from bacterial and mammalian genomes, we found that starting from tetramers, the specificity of VCs was sufficient to detect true read overlaps with high precision. We thus used tetramers, which generates 136-dimensional data points for the VCs.

With default settings, KD-tree generated 2–3 times fewer overlapping pairs than MHAP (Table 1). Yet, we obtained complete and single-contig assemblies for both the PacBio and ONT *E. coli* long read datasets, using KD-tree together with the Canu assembler (v1.5; Koren *et al*., 2017). This indicates that exhaustive overlap detection is not necessary for genome assembly, especially if the overlapping is precise and sequencing depth high. While this was observed (and exploited) before (Li, 2016), we found that a sensitivity as low as 20% is sufficient for single-contig assembly of microbial genomes. Subsequent assembly polishing, e.g. using Quiver (Chin *et al*., 2013), Nanopolish (Loman *et al*., 2015), or Racon (Vaser *et al*., 2017), is recommended in either case and – as long as the assembly is structurally correct – determines the per-base assembly quality.

**Table 1:** Sensitivity, precision, and F1 scores on real PacBio and ONT long read datasets. We determined the ground truth to compare against by mapping the reads to the finished *E. coli* K12 MG1655 reference genome. KD-tree is less sensitive – by design – but as precise as other tools.

| Software | PacBio P6-C4 *E. coli* |  |  | ONT SQK-MAP-006 *E. coli* |  |  |
| --- | --- | --- | --- | --- | --- | --- |
|  | Sens. (%) | Prec. (%) | F1 (%) | Sens. (%) | Prec. (%) | F1 (%) |
| BLASR* | 66.0 | 96.5 | 78.3 | 89.9 | 73.0 | 80.6 |
| DALIGNER* | 83.8 | 85.8 | 84.8 | 92.9 | 91.0 | 91.9 |
| GraphMap* | 71.7 | 94.0 | 81.4 | 90.6 | 93.4 | 92.0 |
| MHAP | 46.2 | 86.7 | 60.3 | 83.9 | 77.7 | 80.7 |
| MHAP* | 79.8 | 79.8 | 79.8 | 91.2 | 82.0 | 86.3 |
| Minimap | 74.3 | 94.2 | 83.0 | 91.1 | 98.0 | 94.4 |
| KD-tree | 19.7 | 80.1 | 31.6 | 43.7 | 86.9 | 58.1 |

\*Values taken from Chu *et al.*, 2017, who derived these from the best settings of each tool (according to the highest F1 score) after an exhaustive parameter search. We include results for the default settings of MHAP and the recommended *ava10k* preset for PacBio/ONT all-vs-all read mapping of Minimap for comparison.

For a region with coverage *N*, the number of all possible overlap pairs is (*N −* 1)^2^*/*2, while the number of non-redundant overlaps is only (*N −* 1). A quadratic growth of runtime for high coverage datasets is avoided by our algorithm by reporting a limited number of ANNs (40) for each tag.

### 3.2 Computational performance

We measured the runtime and memory usage for KD-tree, MHAP, and Minimap. MHAP is the default and widely used overlapper in the Canu assembly pipeline and Minimap is the most computationally efficient software to date (Chu *et al*., 2017). All programs were run on a single CPU, though the FLANN software library supports and scales well also to multiple cores and distributed configurations.

KD-tree was faster than MHAP and required less memory than MHAP and Minimap (Table 2). For the ONT *E. coli* dataset with ~40,000 reads, loading, embedding, and tagging required 12 minutes. The tag number was ~625,000, the indexing time one minute, and the ANN search time less than two minutes, amounting to a total runtime of 18 minutes. In comparison, MHAP required one hour to detect read-to-read overlaps for this dataset. We anticipate that with less sensitive settings (suitable for higher coverage and lower noise) for a PacBio read dataset, the speed of MHAP would improve.

**Table 2:** Runtime and memory usage on real long read datasets. The benchmarking PacBio and ONT datasets were generated from *E. coli* K12 MG1655, and contain 82,738 reads (748 Mbp) and 39,921 reads (388 Mbp), respectively. Elapsed time and peak memory usage were measured using GNU time for single-threaded program execution.

| Software | PacBio P6-C4 *E. coli* |  | ONT SQK-MAP-006 *E. coli* |  |
| --- | --- | --- | --- | --- |
|  | Elapsed time<br>(h:mm:ss) | Peak memory<br>usage (GB) | Elapsed time<br>(h:mm:ss) | Peak memory<br>usage (GB) |
| MHAP | 1:54:35 | 28.96 | 1:00:06 | 19.33 |
| Minimap | 0:11:13 | 6.55 | 0:06:25 | 3.30 |
| KD-tree | 0:37:04 | 2.01 | 0:18:25 | 1.06 |

KD-tree also demonstrated a substantial decrease in memory consumption; it required 15–20 times less memory than MHAP (and a few times less than Minimap) for these datasets. Aside from the implementation specifics we assume that the memory efficiency of our implementation is related to the higher compactness of the KD-tree index in comparison to the hash tables used by other programs (Muja and Lowe, 2014).

## 4 Discussion

We have demonstrated the value of a new kind of algorithm for overlapping long and noisy reads, which uses short *k*-mer spectra and ANN KD-tree search. Our implementation performed comparably and in some aspects (e.g. memory efficiency, algorithmic and software implementation simplicity) challenging to state-of-the-art overlappers.

The context-aware tagging technique implemented in KD-tree limits the growth of the index size with increasing coverage, thus allowing a more efficient processing of high-coverage datasets. The computational complexity for all-vs-all comparisons of *N* data points with KD-trees is still *N* * *log*(*N*), similar to other overlappers. With context-unaware indexing of a read dataset, the index size will grow linearly with the dataset size. However, if the index is constructed dynamically, discarding redundant entries on-the-fly, our tagging technique can allow processing time to grow linearly with data set size, as the index size stops growing with coverage. Specifically, for high-coverage (hence high redundancy) datasets, the dictionary size of unique (context-specific) tags (and hence the index) can be limited in a genome as coverage grows to multiples of genome size. For example, tagging a mammalian genome with one tag per 1 kb average density would require about 3 millions unique tags, which require less than 4 GB of memory in a KD-tree index. Sequencing data can then be streamed efficiently through this index without further memory consumption (since the index is complete), and each read will be aligned to a few tags, producing an overlap graph.

Surprisingly, short *k*-mers, starting from tetramers we used here, already produced sufficiently precise spectrums for finding overlaps. In comparison, other overlappers and aligners use relatively large values of *k* (>10). The algorithm performed well also for short tags (i.e. from 50 bp), thus another usage could be the joint processing of hybrid data sets with both short and long reads (Koren *et al*., 2012; Kunath *et al*., 2017). Similarly to Minimap and other programs, it would also be applicable for whole-genome alignments and comparisons, and possibly for other -omics applications where structuring of multidimensional data is needed.

Likely further gains in speed and memory efficiency are possible. For example, it might be possible to significantly reduce the considered number of *k*-mers by with a minimizer-like approach (Roberts *et al*., 2004). We also found that on-the-fly points insertion/deletion into the KD-tree index and searching tags one-by-one (not in a single batch) did not significantly affect performance, so one could build an index discarding redundant tags one-the-fly, further improving efficiency for datasets with high coverage.

With read lengths and throughput of the long reads sequencing technologies continuously increasing at simultaneously decreasing costs, there will be emerging demand for methods allowing efficient handling of long reads. Single-molecule technologies are likely to retain substantial noise, inherent to experimenting at this physical scale. Hence the demand for efficient error-tolerant overlappers and aligners for long reads is likely sustainable. At the same time the trend of miniaturization of sequencers (e.g. ONT’s MinION) and their use as portable devices in field applications (Quick *et al*., 2016; Faria *et al*., 2017) will further increase the need for computational processing efficiency.

## Acknowledgements

We thank Adrian Fritz for feedback on earlier versions of the manuscript, Justin Chu for helpful advice about benchmarking data and setup, and Peter Belmann for building the KD-tree biobox.

## Funding

Supported by DZIF (German Center for Infection Research).

## References

Belmann, P. et al. (2015). Bioboxes: standardised containers for interchangeable bioinformatics software. Gigascience, 4, 47. doi:10.1186/s13742-015-0087-0.

Bentley, J. L. (1975). Multidimensional binary search trees used for associative searching. Commun. ACM, 18(9), 509–517. doi:10.1145/361002.361007.

Berlin, K. et al. (2015). Assembling large genomes with single-molecule sequencing and locality-sensitive hashing. Nat. Biotechnol., 33(6), 623–630. doi:10.1038/nbt.3238.

Chin, C. S. et al. (2013). Nonhybrid, finished microbial genome assemblies from long-read SMRT sequencing data. Nat. Methods, 10(6), 563–569. doi: 10.1038/nmeth.2474.

Chu, J. et al. (2017). Innovations and challenges in detecting long read overlaps: an evaluation of the state-of-the-art. Bioinformatics, 33(8), 1261–1270. doi:10.1093/bioinformatics/btw811.

Faria, N. R. et al. (2017). Establishment and cryptic transmission of Zika virus in Brazil and the Americas. Nature, 546(7658), 406–410. doi:10.1038/nature22401.

Goodwin, S. et al. (2016). Coming of age: ten years of next-generation sequencing technologies. Nat. Rev. Genet., 17(6), 333–351. doi:10.1038/nrg.2016.49.

Koren, S. et al. (2012). Hybrid error correction and de novo assembly of single-molecule sequencing reads. Nat. Biotechnol., 30(7), 693–700. doi:10.1038/nbt.2280.

Koren, S. et al. (2017). Canu: scalable and accurate long-read assembly via adaptive k-mer weighting and repeat separation. Genome Res., 27(5), 722–736. doi:10.1101/gr.215087.116.

Kunath, B. J. et al. (2017). Metagenomics and CAZyme Discovery. Methods Mol. Biol., 1588, 255–277. doi:10.1007/978-1-4939-6899-2_20.

Laehnemann, D. et al. (2016). Denoising DNA deep sequencing data-high-throughput sequencing errors and their correction. Brief. Bioinformatics, 17(1), 154–179. doi:10.1093/bib/bbv029.

Li, H. (2016). Minimap and miniasm: fast mapping and de novo assembly for noisy long sequences. Bioinformatics, 32(14), 2103–2110. doi:10.1093/bioinformatics/btw152.

Loman, N. J. et al. (2015). A complete bacterial genome assembled de novo using only nanopore sequencing data. Nat. Methods, 12(8), 733–735. doi:10.1038/nmeth.3444.

Marimont, R. B. and Shapiro, M. B. (1979). Nearest neighbour searches and the curse of dimensionality. IMA Journal of Applied Mathematics, 24(1), 59–70. doi:10.1093/imamat/24.1.59.

Muja, M. and Lowe, D. G. (2014). Scalable Nearest Neighbor Algorithms for High Dimensional Data. IEEE Trans Pattern Anal Mach Intell, 36(11), 2227–2240. doi:10.1109/TPAMI.2014.2321376.

Myers, G. (2014). Efficient local alignment discovery amongst noisy long reads. In Lecture Notes in Computer Science, pages 52–67. Springer Berlin Heidelberg. doi:10.1007/978-3-662-44753-6_5].

Quick, J. et al. (2016). Real-time, portable genome sequencing for Ebola surveillance. Nature, 530(7589), 228–232. doi:10.1038/nature16996.

Roberts, M. et al. (2004). Reducing storage requirements for biological sequence comparison. Bioinformatics, 20(18), 3363–3369. doi:10.1093/bioinformatics/bth408.

Sović, I. et al. (2016). Fast and sensitive mapping of nanopore sequencing reads with GraphMap. Nat Commun, 7, 11307. doi:10.1038/ncomms11307.

Vaser, R. et al. (2017). Fast and accurate de novo genome assembly from long uncorrected reads. Genome Res., 27(5), 737–746. doi:10.1101/gr.214270.116.

Wibberg, D. et al. (2016). Finished genome sequence and methylome of the cyanide-degrading Pseudomonas pseudoalcaligenes strain CECT5344 as resolved by single-molecule real-time sequencing. J. Biotechnol., 232, 61–68. doi:10.1016/j.jbiotec.2016.04.008.

